# Small molecule inhibition of human cGAS reduces total cGAMP output and cytokine expression in cells

**DOI:** 10.1101/2020.03.30.016535

**Authors:** Caroline Wiser, Byungil Kim, Jessica Vincent, Manuel Ascano

**Author notes:** MATERIALS & CORRESPONDENCE Correspondence and/or requests for materials should be addressed to M.A.

## Abstract

The cGAS-STING pathway is a major mechanism that mammalian cells utilize to detect cytoplasmic dsDNA from incoming viruses, bacteria, or self. CYCLIC GMP-AMP SYNTHASE (cGAS) is the sensor protein that directly binds dsDNAs. cGAS synthesizes cyclic GMP-AMP (cGAMP), which binds to the adaptor STIMULATOR OF INTERFERON GENES (STING), activating an INTERFERON REGULATORY FACTOR 3 (IRF3)-mediated immune response. Constitutive activation can result in interferonopathies such as Aicardi-Goutieres Syndrome (AGS) or other lupus-like autoimmune disorders. While inhibitors targeting mouse or human cGAS have been reported, the identification of a small molecule that targets both homologs of cGAS has been challenging. Here, we show that RU.521 is capable of potently and selectively inhibiting mouse and human cGAS in cell lines and human primary cells. This inhibitory activity requires the presence of cGAS, but it cannot suppress an immune response in cells activated by RNA, Toll-like receptor ligands, cGAMP, or recombinant interferon. Importantly, when RU.521 is applied to cells, the production of dsDNA-induced intracellular cGAMP is suppressed in a dose-dependent manner. Our work validates the use of RU.521 for probing DNA-induced innate immune responses and underscores its potential as an ideal scaffold towards pre-clinical development, given its potency against human and mouse cGAS.

## INTRODUCTION

The deployment of pattern recognition receptors (PRRs) is the primary method for the innate immune system to detect pathogens or cellular damage via their associated molecular patterns (PAMPs and DAMPs)^1^. Among PRRs are cytoplasmic RNA and DNA nucleic acid sensors^2-9^. Nucleic acids from incoming pathogens have distinct molecular features that are recognized by host PRRs localized within the cytosol^10^. Human CYCLIC GMP-AMP SYNTHASE (h-cGAS)^11,12^ produces the cyclic dinucleotide c[G(2’,5’)pA(3’,5’)p], or cGAMP, upon binding cytoplasmic dsDNA^13,14,15^. cGAMP then binds to the endoplasmic reticulum-bound adaptor protein STIMULATOR OF INTERFERON GENES (STING), and induces a signaling cascade that ultimately results in the phosphorylation, dimerization, and translocation of INTERFERON REGULATORY FACTOR 3 (IRF3) into the nucleus^16,17,18^. The nuclear translocation of activated IRF3 leads to the transcriptional upregulation of type I interferons and pro-inflammatory cytokines^2,19,20^.

An essential role for cGAS is its capacity to detect DNA derived from invading pathogens. Numerous reports have demonstrated the role of cGAS against prokaryotes including *L. monocytogenes*^21^, *C. trachomatis*^22^, *L. pneumophila*^23^, and *M. tuberculosis*^24,25^. Moreover, cGAS has also been reported to detect DNA derived from viruses^26^. Upon infection, the cGAS-STING pathway is activated by viruses, including some members of the herpes family^27^, oncogenic viruses like HPV^28^, and even retroviruses such as HIV^29^. By serving as the first line of defense against a diverse body of pathogenic DNAs, cGAS establishes its necessity to the cell’s innate immune system.

The proper regulation of cGAS is critical to maintain immune homeostasis. The development of patient serum immunoreactivity against dsDNA is a common symptom among patients with systemic lupus erythematosis (SLE)30,31. TREX1 is an exonuclease located in the cytosol that degrades mislocalized DNA^32,33^. However, inactivating mutations in TREX1 can cause prolonged activation of cGAS, resulting in protracted inflammation^34,32,35^. Autoimmune diseases of this nature include Aicardi-Goutières syndrome^36^ and Chilblain lupus^37^, where the overproduction of interferons leads to chronic inflammation. Knockout studies have shown that Trex1 null mice, which have elevated interferon levels, return to WT levels of interferon upon knockout of cGAS^38,39^, STING^40,41^, or IRF3^40^. These studies further validate the importance of cGAS and its role in regulating immunity, and it has become a prime candidate for the development of small molecules that activate or inhibit its immunopotent activity^42,43,44,45^. Identifying an effective cGAS inhibitor could serve as a tool to help assess the contribution of the cGAS-STING signaling axis in various immune responses, and could ultimately serve as a chemical scaffold towards the development of clinical compounds for the treatment of associated autoimmune diseases.

We previously identified RU.521 as a compound with inhibitory activity on mouse cGAS^46^. Crystal structures show that RU.521 occupies the catalytic pocket of murine cGAS, thus interfering with the entry of its ATP and GTP substrates. However, questions have arisen regarding the ability of RU.521 to inhibit human cGAS using bacterially expressed proteins^47,44^. Here, we report our results in which we evaluated the effectiveness of RU.521 inhibition of human cGAS in various cellular contexts. We determined the potency of RU.521 against human cGAS by measuring its inhibitory dose response in mouse and human cells, and find that it has a similar IC_50_ as that observed for the mouse homolog. We demonstrate that RU.521 inhibits cytoplasmic DNA-dependent upregulation of IRF3-dependent transcriptional targets only in the presence of an intact cGAS-STING pathway. Next, we exposed THP-1 cells to various PAMPs while in the presence of RU.521, and demonstrate that there are no observable off-target effects – indicating that RU.521 only suppresses innate immune activation induced by dsDNA if cGAS is present. Using tandem LC/MS-MS, we measured intracellular levels of cGAMP produced by human cGAS and find that RU.521 can suppress the production of the second messenger in a dose-dependent manner. Finally, we show that RU.521 can suppress type I interferon expression and protein production in human primary cells. Altogether, our data illustrate the utility of RU.521 in inhibiting both mouse and human cGAS and its potential as a molecular scaffold to be further developed for eventual clinical use.

## RESULTS

### Inhibition of human cGAS-dependent immune activation by RU.521

THP-1 cells are a monocyte-like human cell line that expresses cGAS and STING, which are required for sensing cytoplasmic dsDNA and eliciting IRF3-mediated *IFNB1* upregulation. In THP-1 KO-cGAS cells, introduction of dsDNA does not stimulate *INTERFERON BETA 1* (*IFNB1*) expression^48^ (see also Fig. 3a); the exogenous addition of h-cGAS is necessary for adequate sensing of cytoplasmic dsDNA and IRF3-mediated *IFNB1* signaling^49,14,11^. To test whether RU.521 could inhibit the human homolog of cGAS, we transfected a plasmid containing h-cGAS into THP-1 KO-cGAS cells and measured IFNB1 mRNA levels in the presence or absence of the inhibitor by RT-qPCR analysis (Fig. 1a, *left panel*). The expression of h-cGAS increased IFNB1 mRNA levels by ∼ 5-fold (4.6x) as compared to untreated cells. Upon addition of RU.521, *IFNB1* expression decreased significantly. We also found that RU.521 could inhibit endogenous h-cGAS in WT (wild type) THP-1 cells that were stimulated with herring testis DNA (HT-DNA) (Fig. 1a *right panel* and Fig 1b) (see also Supplementary Fig. 1 for determination of EC_50_ and EC_90_ values of HT-DNA)^11,50^. Next, we established an RU.521 dose-response curve to determine its IC_50_ (half-maximal inhibitory concentration) for h-cGAS in HT-DNA-activated THP-1 cells (Fig. 1c). Using concentrations spanning the range from 0.001 µM to 100 µM, the IC_50_ was determined to be ∼ 0.8 µM in THP-1 cells. These findings are in line with our previous assessment of RU.521 potency against mouse cGAS (m-cGAS), which was measured to be 0.7 µM in murine RAW 264.7 cell^46^. These results indicate that RU.521 is capable of inhibiting DNA- and h-cGAS-dependent immune activation.

**Figure 1.**
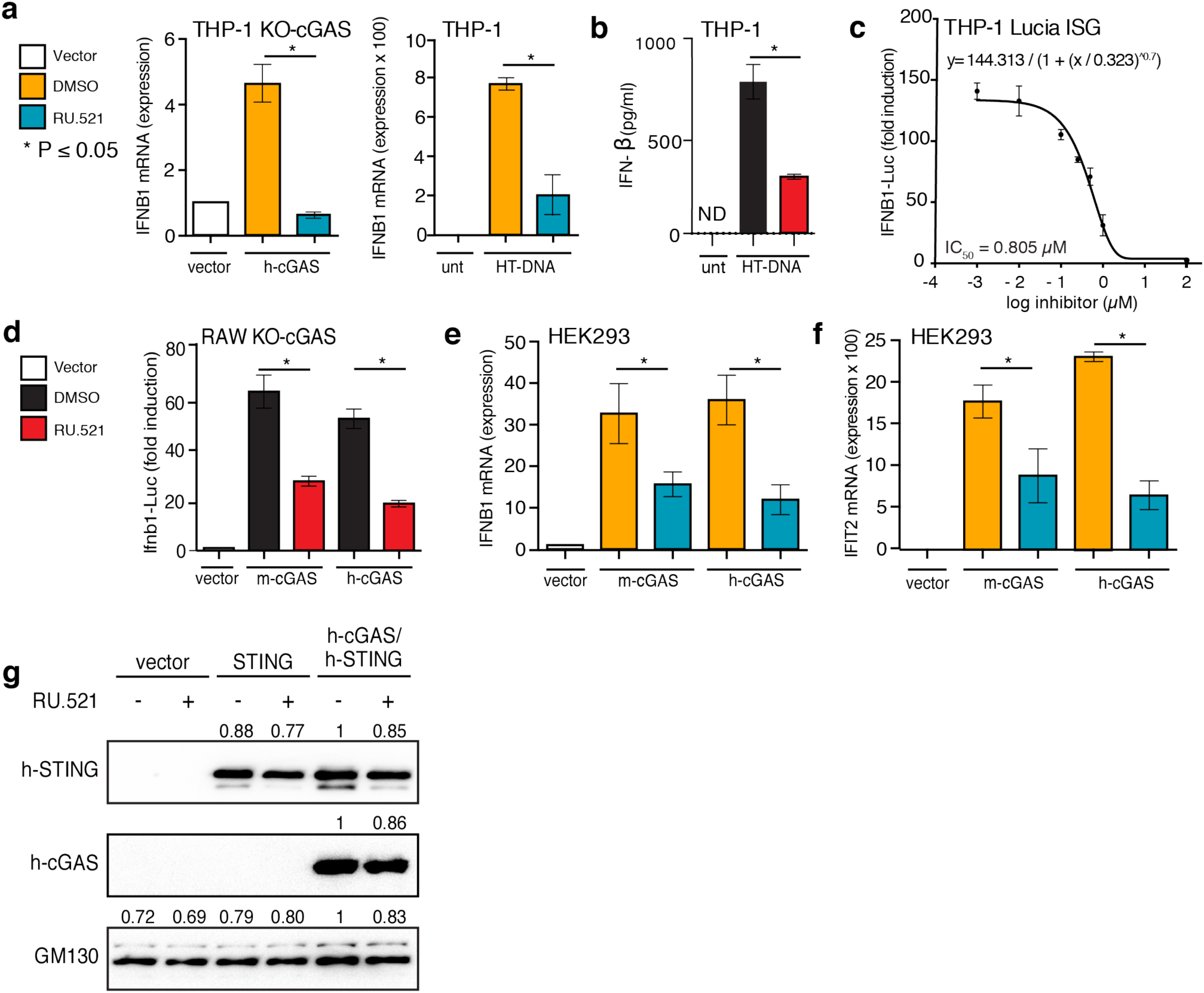
Inhibition of human and mouse cGAS with RU.521 reduces interferon expression in multiple cell types. Lines used were: THP-1 and its derivatives (**a-c**), RAW 267.4 (**d**), and HEK293 (**e-g**) cells. THP-1 KO-cGAS cells were transfected with either vector plasmid control or a plasmid containing h-cGAS, and treated with either vehicle (DMSO) or RU.521 (**a**, *left panel*). WT THP-1 cells were mock treated (unt) or stimulated with 0.3 µg/mL HT-DNA +/- RU.521, followed by RT-qPCR analysis for IFNB1 expression (**a**, *right panel*). An ELISA for human IFNB1 was also performed on THP-1 cells stimulated with HT-DNA +/- RU.521 (0.8 µM) (**b**). A dose-response curve was generated for RU.521 using immune-activated THP-1 Lucia ISG cells (**c**). RAW-Lucia ISG KO cGAS cells were transfected with plasmids expressing WT m-cGAS or h-cGAS, and activity was read via luciferase reporter (**d**). HEK293 cells were transfected with human or mouse cGAS in addition to h-STING, in the presence or absence of RU.521. The RNA levels for IFNB1 and IFIT2 were then measured by RT-qPCR (**e** and **f**). Transfection of vector backbone was used to normalize the data. Immunoblot analysis of transfected h-STING, h-cGAS and m-cGAS into HEK293 cells +/- RU.521 shows protein expression is unaltered by addition of the small molecule. GM130 was used to normalize the data (**g**). The minimum numbers of biological and technical replicates for the assays were three and three, respectively (n = 9). *Error bars* represent SEM. Asterisks (*) denote P ≤ 0.05 where an unpaired t-test with Welch’s correction was used to compare the results.

### Comparison of human and mouse cGAS in response to RU.521

In order to make a more direct comparison of RU.521’s potency between human and mouse cGAS, we expressed either homolog in RAW 264.7 cGAS-knockout macrophages (RAW KO-cGAS), which contain a luciferase reporter gene (ISG54) sensitive to type-I interferon activation. As with THP-1 cells, the absence of cGAS renders RAW cells incapable of responding to exogenous DNA (Fig. 1d). However, transfection of a plasmid containing h-cGAS increased IFNB1-Luc activity by ∼55-fold (55.9x) as compared to vector control. Similarly, expression of mouse cGAS increased IFNB1-Luc activity by ∼60 fold (58.8x). Upon addition of RU.521 (0.8 µM) in cells, which were also transfected with either human or mouse cGAS, we observed a significant reduction in luciferase activity (Fig. 1d).

To demonstrate that RU.521 can inhibit DNA-dependent cGAS activity in other cellular contexts, we measured induction of *IFNB1* and *IFIT2* in HEK293 cells that were co-transfected with human STING (h-STING) and human or mouse cGAS (Fig. 1e-f). When treated with RU.521, *IFNB1* expression was reduced by approximately half for mouse and human cGAS. A similar result was seen with *IFIT2*. We did not observe that RU.521 destabilized HT-DNA *in vitro* (Supplementary Fig. 1b). The protein expressions of transfected h-cGAS, m-cGAS, and h-STING were confirmed to be similar and unaffected by the presence of RU.521 (Fig. 1g and Supplementary Fig. 1c and 2).

### RU.521 selectively inhibits the human cytoplasmic DNA sensing pathway

We next wanted to assess whether RU.521 specifically inhibits the human cytoplasmic DNA sensing pathway via its action on h-cGAS. We therefore exposed THP-1 cells to different ligands of various PRRs (5’ ppp-HP20 (RIG-I like receptors, RLRs), Pam3CSK4 (Toll like receptor, TLR 1/2), poly(I:C) (TLR 3), and lipopolysaccharide (LPS) (TLR 4)), in the presence or absence of RU.521, and measured the levels of IL6 and IFNB1 mRNA by RT-qPCR analysis (Fig. 2 and ^46^). To ensure that the RU.521 compound was functional in all assays, we also exposed THP-1 cells to HT-DNA in parallel. Although THP-1 cells were robustly activated by each PAMP (Supplementary Fig. 3a-l), we found that RU.521 could only inhibit HT-DNA-dependent IFNB1 and IL6 expression (Fig. 2a-b). We also did not observe changes in the expression of IL1B or IL18 levels upon exposure to RU.521 (Fig. 2c). The protein levels of cGAS were not altered in the presence of any ligand, or by the addition of RU.521 (Fig. 2d, and Supplementary Fig. 3m), discounting the potential of cGAS protein levels indirectly contributing to changes in the sensitivities of the various PAMPs.

**Figure 2.**
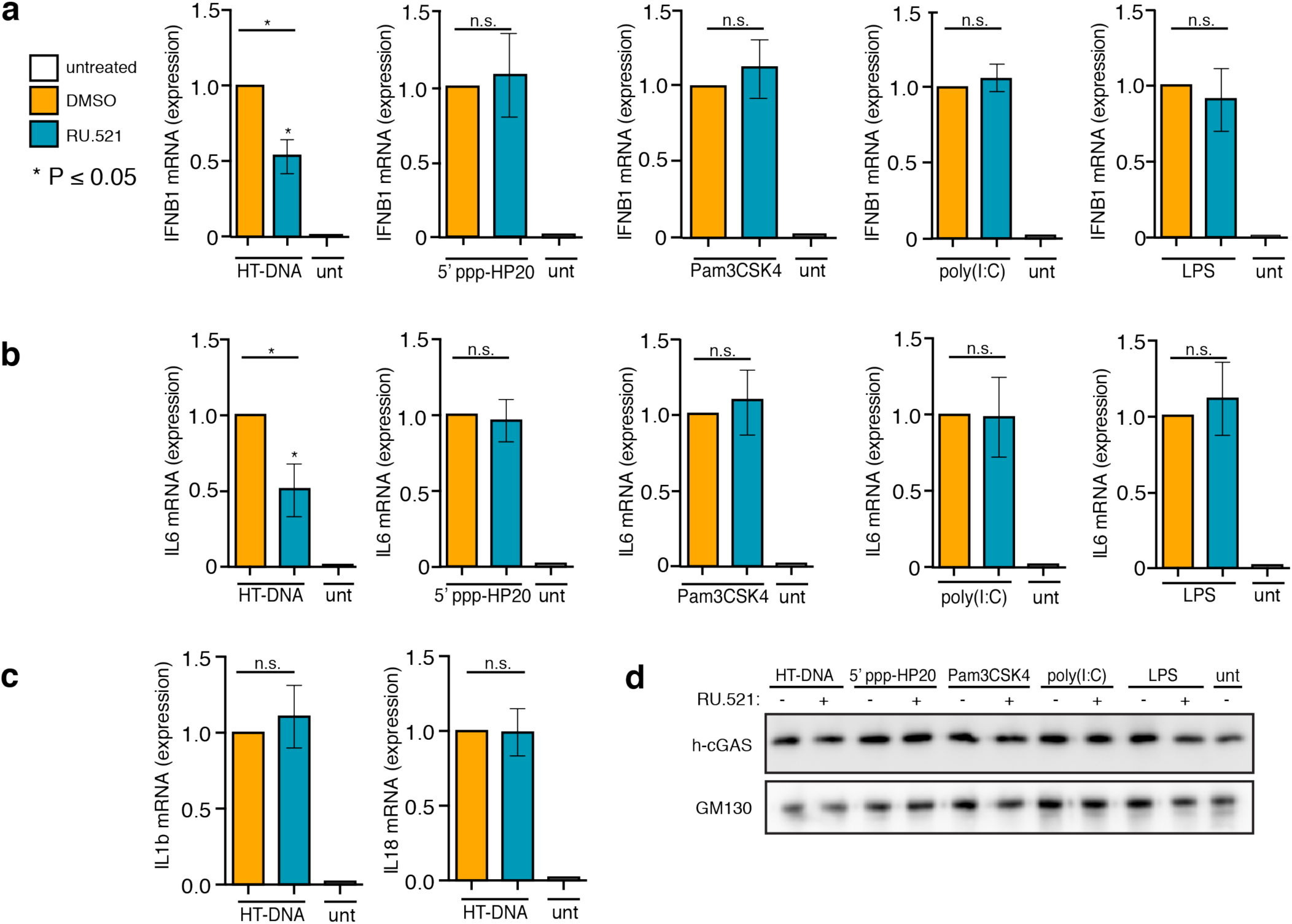
Potent and selective inhibition of h-cGAS activity in THP-1 cells. WT THP-1 cells were exposed to HT-DNA, 5’ppp-HP20 RNA, PAM3CSK4, poly(I:C), lipopolysaccharide (LPS), to promote either a type I interferon (IFNB1) (**a**) or NF-κB (IL6) (**b**) response under different immune stimuli and *simultaneously* treated with 0.8 µM of RU.521. After being treated for 24 h, cells were processed for RT-qPCR analysis. An RT-qPCR analysis of IL1B and IL18 expression was also performed in cells exposed to HT-DNA to determine whether RU.521 might suppress their expression (**c**). Western blot analysis on endogenous h-cGAS was performed to determine whether PRR ligand or RU.521exposure affected its protein levels (**d**). An immunoblot against GM130 served as loading control. The minimum numbers of biological and technical replicates for the assays were two and three, respectively (n = 6). *Error bars* represent SEM. Asterisks (*) denote P ≤ 0.05 where an unpaired t-test with Welch’s correction was used to compare the results of each stimulant to dsDNA.

### RU.521 specifically targets cGAS in the cGAS-STING signaling pathway

To ensure RU.521 specifically inhibited cGAS and not downstream signaling events, we transfected THP-1 KO-cGAS cells with HT-DNA in the presence or absence of RU.521 and measured IFNB1 expression by RT-qPCR analysis (Fig. 3a). As expected, IFNB1 expression remained low and unperturbed in THP-1 KO-cGAS cells exposed to HT-DNA or HT-DNA with RU.52 (Fig. 3a, *left panel*). Cells directly stimulated with chemically synthesized cGAMP resulted in appreciable upregulation of IFNB1, indicating that STING can be activated. However, RU.521 was unable to inhibit cGAMP-mediated IFNB1 induction, as it could in THP-1 cells that were stimulated with DNA - which were performed in a parallel experiment (Fig. 3a, *right panel*). These results demonstrate that RU.521 does not inhibit downstream effectors of cGAS. Given that a number of our reporter assays are based on the use of the type I interferon-sensitive ISG54 promoter, we next tested whether RU.521 would have any inhibitory effects on cells directly stimulated by recombinant IFNB1 and subsequent IFNAR-dependent JAK-STAT pathway activation. We stimulated THP-1 cells with recombinant human IFNB1 in the presence or absence of RU.521 (Fig. 3b). THP-1 cells activated by recombinant IFNB1 were refractory to the addition of RU.521, in contrast to when cells are stimulated with HT-DNA. These results indicate that direct immune activation of THP-1 cells with synthetic cGAMP or recombinant IFNB1 protein is not inhibited by the presence of RU.521. We next measured the cytotoxicity of RU.521 in THP-1 cells through a dose-response analysis (Fig. 3c). The LD_50_ was determined to be 31.4 µM, which is approximately 40 times greater than its IC_50_. These results indicated that the working concentrations of RU.521 used in our study were not a result of potentially confounding toxic effects.

**Figure 3.**
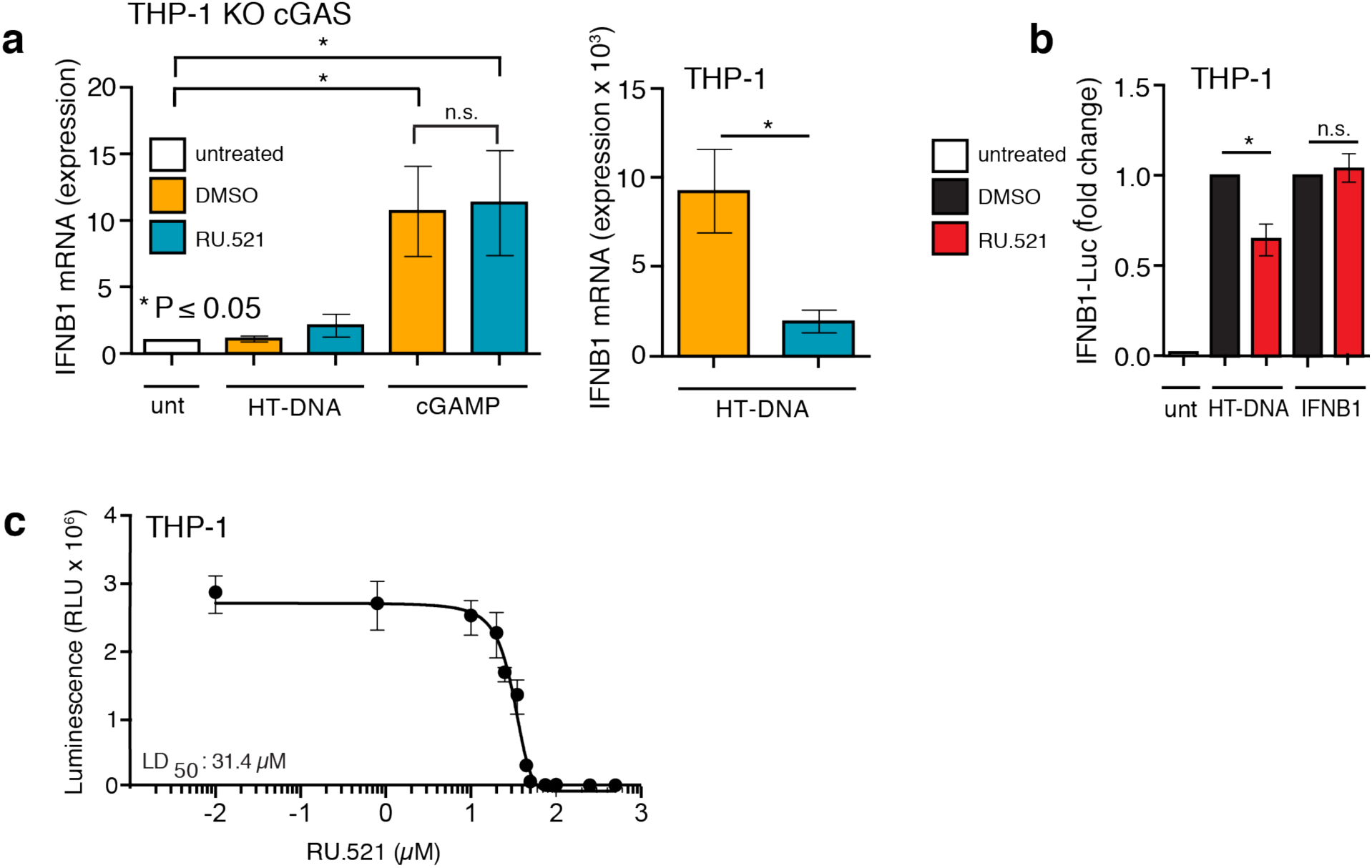
RU.521 acts upstream of cGAMP and downstream of HT-DNA in the cGAS-STING signaling axis. THP-1 KO cGAS cells were treated with HT-DNA or cGAMP in the absence and presence of RU.521 (0.8 µM) (**a**, *left panel*). In a parallel assay, WT THP-1 cells were exposed to HT-DNA +/- RU.521 followed by RT-qPCR analysis for IFNB1 expression (**a**, *right panel*). WT THP-1 Lucia ISG cells were stimulated with HT-DNA or recombinant IFNB1 +/- RU.521, followed by luciferase assay to measure type 1 interferon activation (**b**). A cytotoxicity dose response curve in THP-1 cells was performed across indicated RU.521 concentrations to calculate LD_50_ (LD_50_ = 31.4 µM) (**c**). The minimum numbers of biological and technical replicates for the assays were two and three, respectively (n = 6). *Error bars* represent SEM. Asterisks (*) denote P ≤ 0.05 where an unpaired t-test with Welch’s correction was used to compare the results.

### DNA-stimulated cellular cGAMP levels are reduced in the presence of RU.521

In light of the discrepancies between data observed by others on the *in vitro* analyses of recombinant cGAS with RU.521^51,47,44^ and our results thus far in this report, we sought a more direct measure of cGAS activity in cells. Since cGAMP is the enzymatic product of cGAS activity, quantification of cellular cGAMP levels would provide the most direct readout of RU.521 action inside cells. We therefore sought to measure cGAMP intracellular levels from HT-DNA-activated cells by LC-MS/MS (Fig. 4a, schematic)^52,20^. We transfected RAW KO-cGAS cells with either m-cGAS or h-cGAS and treated with 0 µM, 0.8 µM, or 3 µM RU.521 (control, IC_50_, and IC_90_ respectively) and verified RU.521 dose-dependent activity using the luciferase reporter (Fig. 4b). RAW KO-cGAS cells were chosen since we established that we can heterologously express human or mouse cGAS with similar transfection efficiency, unlike the limited transfection efficiency achievable in THP-1 cells. An LC-MS/MS reference curve was established using synthetic cGAMP (Fig. 4c). From the same cellular lysates used to measure luciferase activity in Fig. 4b, we took aliquots and processed these samples for LC-MS/MS analysis (Fig. 4d). We could not detect basal levels of cGAMP from RAW KO-cGAS cells. Addition and stimulation of m-cGAS resulted in detectable cellular cGAMP levels (∼28.6 femtomoles), which were reduced in samples treated with RU.521. Addition and stimulation of h-cGAS led to lower cGAMP levels (7.3 femtomoles) in cells, whose cyclase activity is known to be less catalytic than its murine counterpart^13,46,47^. Nonetheless, we found that increasing doses of RU.521 reduced cellular cGAMP levels synthesized by h-cGAS accordingly.

**Figure 4.**
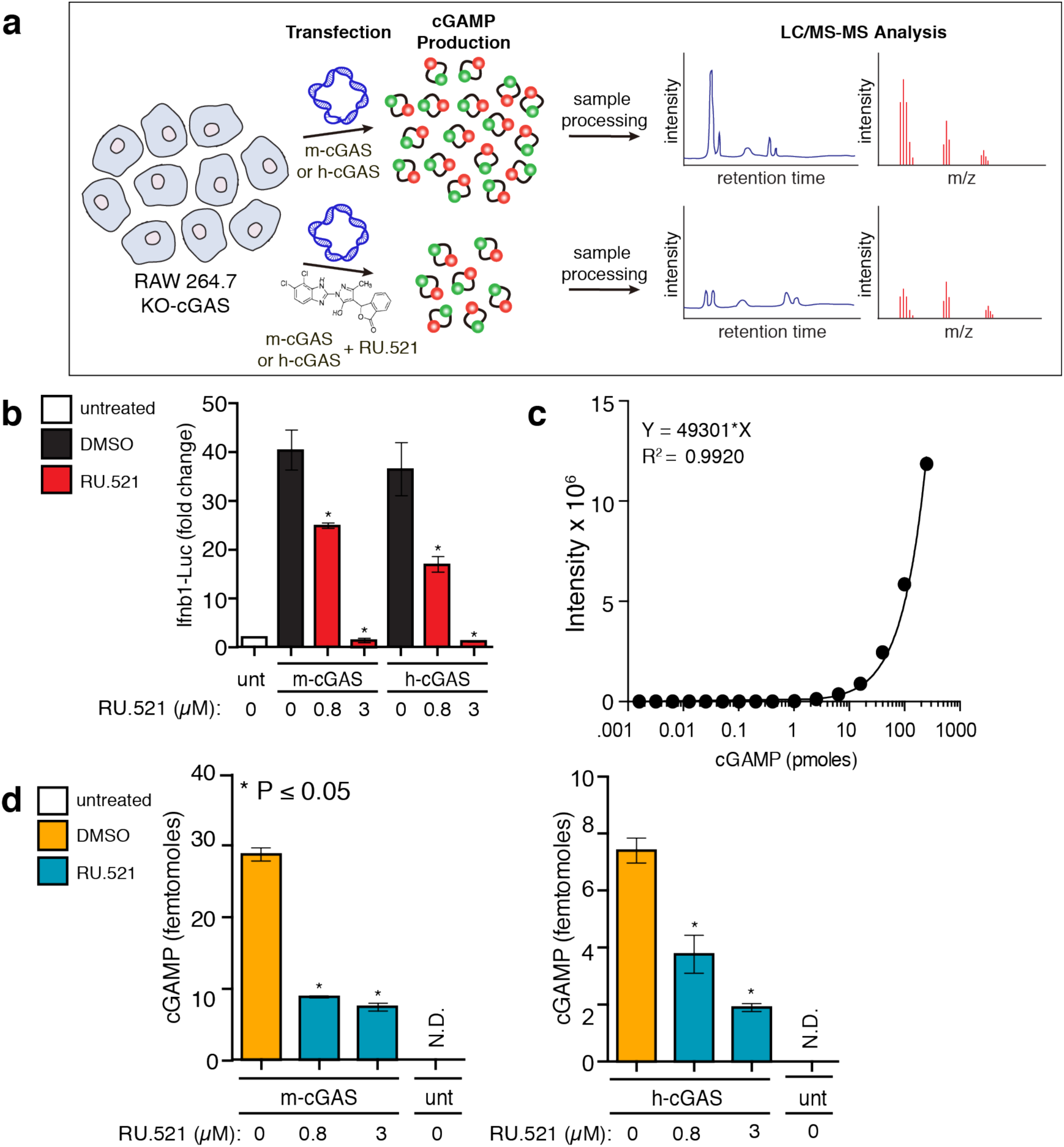
RU.521 reduces intracellular cGAMP levels produced by human and mouse cGAS. RAW-Lucia ISG-KO-cGAS cells were transfected with m-cGAS or h-cGAS and exposed to RU.521. After 24 h incubation, cells were collected and processed for luciferase assays and LC-MS/MS analysis (**a**). Luciferase activity was monitored to validate RU.521 effects on interferon induction, as before (**b**). The intracellular levels of cGAMP were measured by LC-MS/MS (**c, d**). A reference curve was established using synthetic cGAMP (**c**). Femtomole quantities of cGAMP were detected in samples generated by mouse or human cGAS activity in cells, in the presence or absence of RU.521 treatment (**d**). The minimum numbers of biological and technical replicates for the assays were three and three, respectively (n = 9). *Error bars* represent SEM. Asterisks (*) denote P ≤ 0.05 where an unpaired t-test with Welch’s correction was used to compare the results.

### RU.521 shuts down DNA-dependent interferon activation in human primary cells

Finally, we tested whether RU.521 could suppress type I interferon expression in human primary peripheral blood mononuclear cells (PBMCs) and M1-macrophage differentiated primary cells (Fig 5). Blood was taken from three healthy donors, from which PBMCs were isolated or further differentiated into M1 macrophages. PBMCs were transfected with HT-DNA in the absence or presence of increasing RU.521 (IC_50_ and IC_90_), followed by RT-qPCR analysis (Fig. 5a). M1 macrophages were similarly exposed to HT-DNA and treated with RU.521 (IC_90_ only), followed by ELISA to measure secreted IFNB1 protein levels (Fig. 5b). In both cases, we see that cells treated with RU.521 effectively suppressed HT-DNA mediated interferon activation.

**Figure 5.**
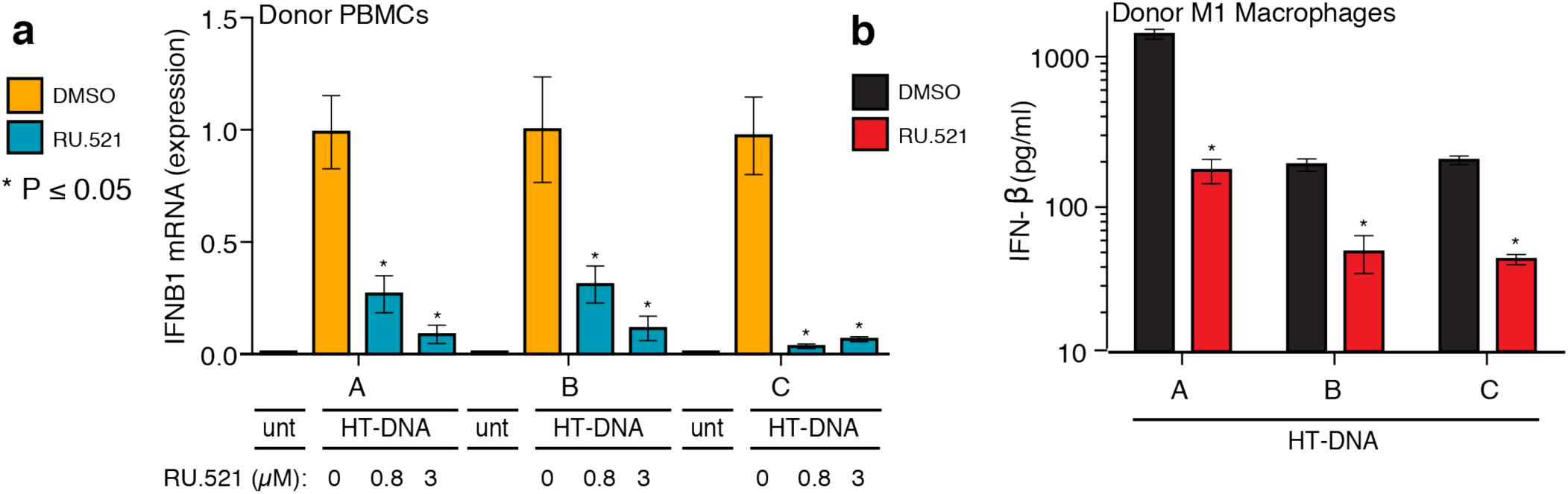
RU.521 suppresses cGAS activity in human primary cells from healthy donors. PBMCs were isolated from three human donors and and either directly stimulated with HT-DNA (**a**) or further differentiated into M1 macrophages (**b**). RU.521 was also added to a subset of the PBMCs at IC_50_ and IC_90_ (0.8 µM and 3 µM, respectively), followed by RT-qPCR analysis for IFNB1 expression (**a**). M1 macrophages were exposed to HT-DNA +/- RU.521, followed by ELISA for secreted IFNB1 protein levels (**b**). For PBMCs, each donor had (n = 9) technical replicates; for M1 macrophages, each donor had (n = 4) technical replicates. *Error bars* represent SEM. Asterisks (*) denote P ≤ 0.05 where an unpaired t-test with Welch’s correction was used to compare the results.

## DISCUSSION

Given the importance of the cGAS pathway to stimulating the innate immune system, a number of agonists and antagonists have been generated in attempts to probe the molecular underpinnings and to manipulate activity for therapeutic intervention^53,43,54,45,44^. While *in vitro* studies have difficulties in showing that RU.521 can inhibit recombinant human cGAS^47,44^, our data here indicate that RU.521 does inhibit human cGAS activity in cells. We speculate that the discrepancies may reside with the differences between purifying recombinant human protein and *in vitro* experimentation, versus assaying endogenous or mammalian-expressed cGAS where post-translational modifications and subcellular localization of this protein are known to play essential roles to its regulation^55,56,50,57^. We specifically took advantage of well-characterized mouse and human cell lines for which the state of cytoplasmic DNA sensing is best understood - and for which it is clear that in the absence of cGAS and/or STING, DNA- and IRF3-dependent transcriptional responses cannot be upregulated. Exposure of THP-1 KO-cGAS or RAW KO-cGAS cells to DNA does not elicit an immune response unless exogenous cGAS is provided. For HEK293 cells, both cGAS and STING are required. Using THP-1, HEK293 and RAW264.7 cells, we directly compared the inhibitory effects of RU.521 against human and mouse cGAS and found that it had similar potency against either homolog in each cellular context. If RU.521’s inhibitory activity is even partially due to off-target effects, then these same off-targets would have to be conserved among the tested human and mouse cell lines in order to explain our results. Moreover, since the innate immune system is comprised of a network of various signaling pathways, we wanted to ensure that RU.521 was specific only to the cGAS-STING axis. Therefore, we treated WT THP-1 cells with ligands that stimulate various PRRs, added RU.521 to the cells, and then measured *IL6* and *IFNB1* mRNA expression. We observed no inhibition of *IL6* or *IFNB1* transcriptional upregulation upon addition of RU.521 despite stimulating RLRs, TLR1/2, TLR3, and TLR4 with their respective. Only in the presence of HT-DNA did we observe potent inhibition of human interferon sensitive genes, demonstrating RU.521 selectivity in suppressing the cGAS-STING DNA sensing pathway.

In order to confirm what step of the DNA sensing pathway was inhibited by RU.521, we activated cells in two different ways. First we stimulated cells directly with synthetic cGAMP, obviating the need to activate cGAS. And second, we exposed cells to recombinant human IFNB1 protein since the interferon-based reporter or transcriptional assays we had used could be acted upon by RU.521, independent of cGAS. Under these conditions, STING and the full complement of the JAK/STAT pathway are functional. In all cases, cGAS itself is not being stimulated or is absent. We find that RU.521 inhibitory activity in cells is only measurable downstream of cGAS activation by DNA, but upstream of cGAMP or IFNB1 stimulation.

We then used mass spectrometry to quantify the intracellular levels of cGAMP produced by cGAS. Given the difficulties reported by others with purely *in vitro* systems, we reasoned that measuring cellular cGAMP would be an ideal alternative for assessing RU.521 on cGAS. We used RAW KO-cGAS cells as a background to express either human or mouse cGAS and found that RU.521 could reduce the cGAMP levels of human or mouse cGAS in a dose-dependent fashion. Importantly, the lysates from these experiments were orthogonally corroborated using a luciferase assay. Finally, we used human primary cells to demonstrate that RU.521 is as potent in suppressing DNA-induced interferon activation as that observed in the cell lines used in this study.

Taken together, our results show that RU.521 can inhibit the dsDNA-dependent enzyme activity of human and mouse cGAS, making it the only reported molecule that can inhibit both homologs. We have demonstrated that RU.521 has similar potency and selectivity for either cGAS homolog, independent of the cellular contexts we used. Further, RU.521 can effectively interfere with the production of intracellular cGAMP. These findings support the use of RU.521 in studies interested in inhibiting the cytoplasmic DNA sensing pathway, and as a molecular scaffold for further derivatization towards clinical development.

## METHODS

### Cell lines

Cell lines were cultured and maintained as previously described^46^. Briefly, mouse RAW macrophages (RAW-Lucia ISG, RAW-Lucia ISG-KO-cGAS, Invivogen) were cultured in DMEM (Thermo Fisher cat. 11965) supplemented with 10% FBS, 100 U ml–1 penicillin-streptomycin, Normocin (100 µg ml–1, Invivogen), and Zeocin (200 µg ml–1, Invitrogen) and an additional 20 mM L-glutamine and 1 mM sodium pyruvate. RAW-Lucia cells stably expressed an interferon sensitive response element from the mouse Isg54 minimal promoter and five interferon-stimulated response elements (Isre-Isg54) coupled to a synthetic coelenterazine-utilizing luciferase. Human THP-1 monocyte-like cells (THP-1 Lucia ISG, Invivogen) were maintained in RPMI 1640(Thermo Fisher cat. 11875093) supplemented with 10% FBS, 10 mM HEPES, 1 mM sodium pyruvate, 100 U ml-1 penicillin-streptomycin, Normocin (100 µg ml–1, Invivogen), and Zeocin (200 µg ml–1, Invitrogen) and an additional 2 mM L-glutamine. THP-1-Lucia ISG cells stably express an interferon sensitive response element from the human ISG54 minimal reporter and five interferon-stimulated response elements (ISRE-ISG54) coupled to a synthetic coelenterazine-utilizing luciferase. Human WT THP-1 and THP-1 KO cGAS cells^48^ were maintained in RPMI 1640 with L-glutamine and phenol red (Life Technologies) supplemented with 10% FBS, 10 mM HEPES, 1 mM sodium pyruvate, 100 U ml-1 penicillin-streptomycin, Normocin (100 µg ml–1, Invivogen), and an additional 2 mM L-glutamine. HEK293 cells were maintained in DMEM (Thermo Fisher cat. 11965) supplemented with 10% FBS, 100 U ml–1 penicillin-streptomycin, and an additional 20 mM L-glutamine. All cell lines were routinely checked for mycoplasma contamination using MycoAlert (Lonza).

### Cellular Luciferase Assays

Luciferase assays were performed as previously described^46^. A concentration range of HT-DNA (Sigma D6898) from 0.0015625 µg to 0.8 µg was complexed with Lipofectamine LTX (Life Technologies) per the manufacturer’s suggested protocol. Typically, 0.17 µg HT-DNA was used to stimulate 150,000 THP-1 Lucia ISG cells, which equated to 90% of maximal luciferase activity observed. HT-DNA stimulation of THP-1 cells and subsequent luciferase measurement were performed with or without the addition of RU.521 at indicated concentrations for each figure, or with DMSO as vehicle control. For the dose curve experiments, serial dilutions of RU.521 were applied to 1.5 × 10^5^ cells, plated in 24-well dishes, and co-stimulated with 0.17 µg HT-DNA. In subsequent experiments using RU.521 in THP-1 Lucia ISG cells, the IC_50_ was used while being co-stimulated with one of the following: 50 nM 5’ppp-HP20 hairpin RNA (a kind gift from Anna Pyle, Yale Unversity), 10 µM cGAMP, or 100 U/mL human recombinant Ifn-b (R&D systems). Cells were harvested 18 hrs post-transfection, washed with PBS, and lysed in 1x Luciferase Cell Lysis Buffer (Pierce). Luciferase luminescence was recorded with a microplate reader (BioTek Synergy HTX) using QUANTI-Luc Luciferase reagent (Invivogen) and its suggested protocol (20 µL lysate/well of a 96-well plate; microplate reader set with the following parameters: 50 µL injection, end-point measurement with 4 s start time and 0.1 s reading time).

### PRR Selectivity Assays in THP-1 cells

RU.521 was used at 0.8 µM, representing the IC_50_, and THP-1 cells were co-stimulated with either 400 ng/mL PAM3CSK4 (Invivogen), 50 µg/mL poly(I:C) (Sigma-Aldrich) 200 ng/mL LPS (Sigma-Aldrich), 50 nM 5’ppp-HP20 RNA^58^, 10 µM cGAMP, or 100 U/mL human recombinant Ifn-b (R&D systems) at the cell counts and conditions previously described for the cellular luciferase assays. Cells were harvested 18 h post-transfection, washed with PBS, and RNA was isolated using Trizol (Ambion) and chloroform (Sigma-Aldrich). In all, 1-2 µg total RNA was reverse transcribed using SuperScript IV VILO Master Mix with ezDNase Enzyme (Invitrogen) into cDNA. Real-time PCR was carried out with 1X FastSYBR Green Plus Master Mix (Applied Biosystems) and run on an Applied Biosystems StepOne Plus PCR machine. The expression of mRNAs representing human Tubulin A1A, Il-6, and Ifnb1 measured (See supplementary Table X for primer sequences used.) The target C_T_ values were normalized to Tuba1a C_T_ values and used to calculate ΔC_T._ mRNA expression of target genes were then calculated using the ΔΔC_T_ method (2^ΔΔCT^). Expression values were analyzed as described in the method for cellular luciferase assays. Primer sequences used for qRT-PCR were human *TUBA1A* (F 5’-GAGCGTCCAACCTATACTAACC-3’; R 5’-GCAGCAAGCCATGTATTTACC-3’), *IL6* (F 5’-GACAAACAAATTCGGTACATCCTC-3’; R 5’-CTGGCTTGTTCCTCACTACTC-3’), *IFNB1* (F 5’-CTTCTCCACTACAGCTCTTTCC-3’; R 5’-GCCAGGAGGTTCTCAACAAT-3’), *IFIT2* (F 5’ATGAGTGAGAACAATAAGAATTCCTTGGAG-3’; R 5’-TGCTCTCCAAGGAATTCTTATTGTTCTCAC-3’), *IL18* (F 5’-GATAGCCAGCCTAGAGGTATGG -3’; R 5’-AAGAAGTGCTCGTCCTCGTC-3’), *IL1b* (F 5’-GGTGTCCGTAACTCCGATCC -3’; R 5’-GGTGTCCGTAACTCCGATCC -3’), and mouse *Actb1* (F 5’-CCCTAAGGCCAACCGTGAAAAG-3’; R 5’-AGAGGCATACAGGGACAGCA-3’), *Ifnb1* (F 5’-GAGTTACACTGCCTTTGCCATCC-3’; R 5’-ACTGTCTGCTGGTGGAGTTCAT-3’), *Ifit2* (F 5’-AGTACAACGAGTAAGGAGTCACT-3’; R 5’-CATGTGCAACATACTGGCCT-3’).

### Cytokine expression analysis of THP-1 cells by quantitative PCR

WT and KO-cGAS THP-1 cells were plated at 1.5 × 10^5^ cells/well (24-well dishes), and RU.521 was added to cells at its IC_50_ (0.8 µM). THP-1 and HEK293 cells were then incubated in the presence of RU.521 and harvested roughly 18 hrs later. RNA was extracted using Trizol and chloroform. Genomic DNA was removed used ezDNase (Invitrogen). 1-2 µg of total RNA was reverse-transcribed using oligo d(T)s, random hexamers, and Superscript IV VILO into cDNA. Real-time PCR targeting human *TUBA1A, IL-6, IL-1b, IL-18* and *IFNB1* was carried out and the data was analyzed as previously described in the methods section for cytokine analysis in THP-1 cells.

### Cytotoxicity Assays

RU.521 was serially diluted to concentrations spanning the range tested in the dose-response curve and added to 150,000 THP-1 cells per well in 96 well plates. The cells were then harvested 72 hrs after compound addition. ATP was measured using CellTiter Glo Viability Assay (Promega). Viability values were generated using vehicle (DMSO) or the first dose of RU.521 as 100%.

### HPLC-MS/MS mass spectrometry

RAW KO-cGAS cells were suspended at 1.5 × 10^5^ cells per well; 24 well plates. One plate was untreated, one transfected with mGAS +/- RU.521, and one transfected with h-cGAS +/- RU.521. Two different concentrations of RU.521 were added to the cells: 0.8 µM and 4 µM. Plates were incubated at 37 C / 5% CO_2_ for 24 hours and then collected. Samples were processed as described^52^. Quantum 3 Mass Spec was used with positive ionization to detect cGAMP (retention time 5.30 minutes) in RAW 264.7 KO-cGAS cellular lysates. The final pellets were resuspended in 50 µL H_2_O/ ACN (9:1) with 10 µL injection volume. Each sample was spiked with 7.5 ng Tenofovir (retention time 4.83 min), purchased from Selleck Chemical LLC (S14015MG). Sample analyses were carried out using a Waters Acquity UPLC system (Waters Corp., Milford, MA), made up of a binary solvent manager, refrigerated sample manager, and a heated column manager. Tandem mass spectrometric detection was performed using a TSQ Quantum Ultra triple-stage quadrupole mass spectrometer (Thermo-Electron, San Jose, CA) equipped with a standard ESI ion source. A Waters Atlantis ZIC-cHILIC analytical column (2.1mmc × 150mm, 3 µm analytical column) equipped with a ZIC-cHILIC pre-column was used for all chromatographic separations. Mobile phases were made up of 0.2% acetic acid + 15 mM ammonium acetate in (A) H2O/ACN (9:1) and in (B) H_2_O/ MeOH/ ACN (5:5:90). Gradient conditions were as follows: 0–2 min, B = 85 %; 2-5 min, B = 85-30%; 5–9 min, B = 30 %; 9-11 min, B = 30-85%; 11–20 min, B = 85%. The flow rate was maintained at 300 µl/min in A and software-controlled divert valve was used to transfer eluent from 0-3 min and from 7-20 min of each chromatographic run to waste. The total chromatographic run time was 20 min. The sample injection volume was 10 µl. The mass spectrometer was operated in positive ion mode. Quantitation was based on SRM (Tenofovir: *m/z* 288 → 159, collision energy 27 V, tube lens 100 V; cGAMP: *m/z* 675 → 312, collision energy 24 V, tube lens 177 V). The following optimized parameters were used for the detection of analyte and internal standard: N_2_ sheath gas 20 psi; N_2_ auxiliary gas 5 psi; corona discharge current 10 µA; capillary temperature 300 °C; Ar collision gas 1.0 mtorr; scan time 100 ms; Q3 scan width 0.5 *m/z*; Q1/Q3 peak width at half-maximum 0.7 *m/z*. Data acquisition and quantitative spectral analysis were done using Thermo-Finnigan Xcalibur version 2.0.7 SP1 and Thermo-Finnigan LCQuan version 2.5.6, respectively. Calibration curves were constructed by plotting peak area ratios (Tenofovir/ cGAMP) against analyte concentrations for a series of 10 spiked standards, ranging from 0.0262 picomoles to 250 picomoles cGAMP^52^. A weighting factor of 1/C^2^ was applied in the linear least-squares regression analysis to maintain homogeneity of variance across the concentration range (%RE ≤ 15% at C > LLOQ).

### Human primary cells

Peripheral blood mononuclear cells (PBMC) (kind gift from Rathmell laboratory) isolated from fresh blood of three healthy anonymous donors were prepared from buffy coats using Ficoll-Paque (GE 45-001-749). Monocytes were enriched from PBMCs by Monocyte Attachment Medium (PromoCell 50306290). The monocytes were expanded in culture for 7 d in the presence of 100 ng/ml human GM-CSF. Cells were re-seeded onto 24-well plates at 1 × 10^5^ cells/well and polarized for 48 hr with the addition of 50 ng/ml human IFNγ (R&D systems) for M1 macrophages. These cells were subsequently treated with HT-DNA in the presence of Lipofectamine 2000 (Life Technologies) for an additional 16 hr in the absence of polarizing cytokines.

### Statistical Analyses

The statistical analyses were performed as before^46^. The numbers of biological and technical replicates were three and three, respectively, unless indicated otherwise. Analysis of luminescence values were evaluated for outliers (one standard deviation above and below the mean) for each biological replicate, and the resulting means were used to generate graphs in GraphPad Prism (v. 6.0). Where applicable, an unpaired t-test with Welch’s correction was used to compare the results of each stimulant to dsDNA. The experimental variances observed within control samples (untreated or vehicle) and treated samples were similar.

## ACKNOWLEDGEMENTS

We would like to thank the Vanderbilt Mass Spectrometry Research Center, particularly Dr. Wade Calcutt and James Carmichael, for their assistance with our mass spec experimental design and analysis. We would like to thank members of Dr. Jeff Rathmell’s laboratory, especially Sam Schaefer, for their assistance with PBMC collection. We would like to thank Dr. Plamen Christov and the Vanderbilt Chemical Synthesis Core for their synthesis of RU.521. A special thank you to Dr. Anna Pyle and her laboratory for providing 5’ppp-HP20 RLR ligand. Finally, we would like to thank members of the Ascano laboratory for their insightful advice, assistance, and critical review of the manuscript and project. This work was supported, in part, by the following agencies: 1R35GM119569-04 (M.A.) and 1R01HL145477-01 (Sur PI, M.A. Co-I).

## AUTHOR CONTRIBUTIONS

C.W. contributed to the design, execution, and interpretation of the majority of cellular assays with the assistance of J.V. and B.K. C.W. and M.A. wrote and co-edited the manuscript with assistance from B.K. and the rest of the Ascano laboratory.

