## Supplementary Figures for "Small molecule inhibition of human cGAS reduces total cGAMP output and cytokine expression in cells"

Supplementary Figure 1:

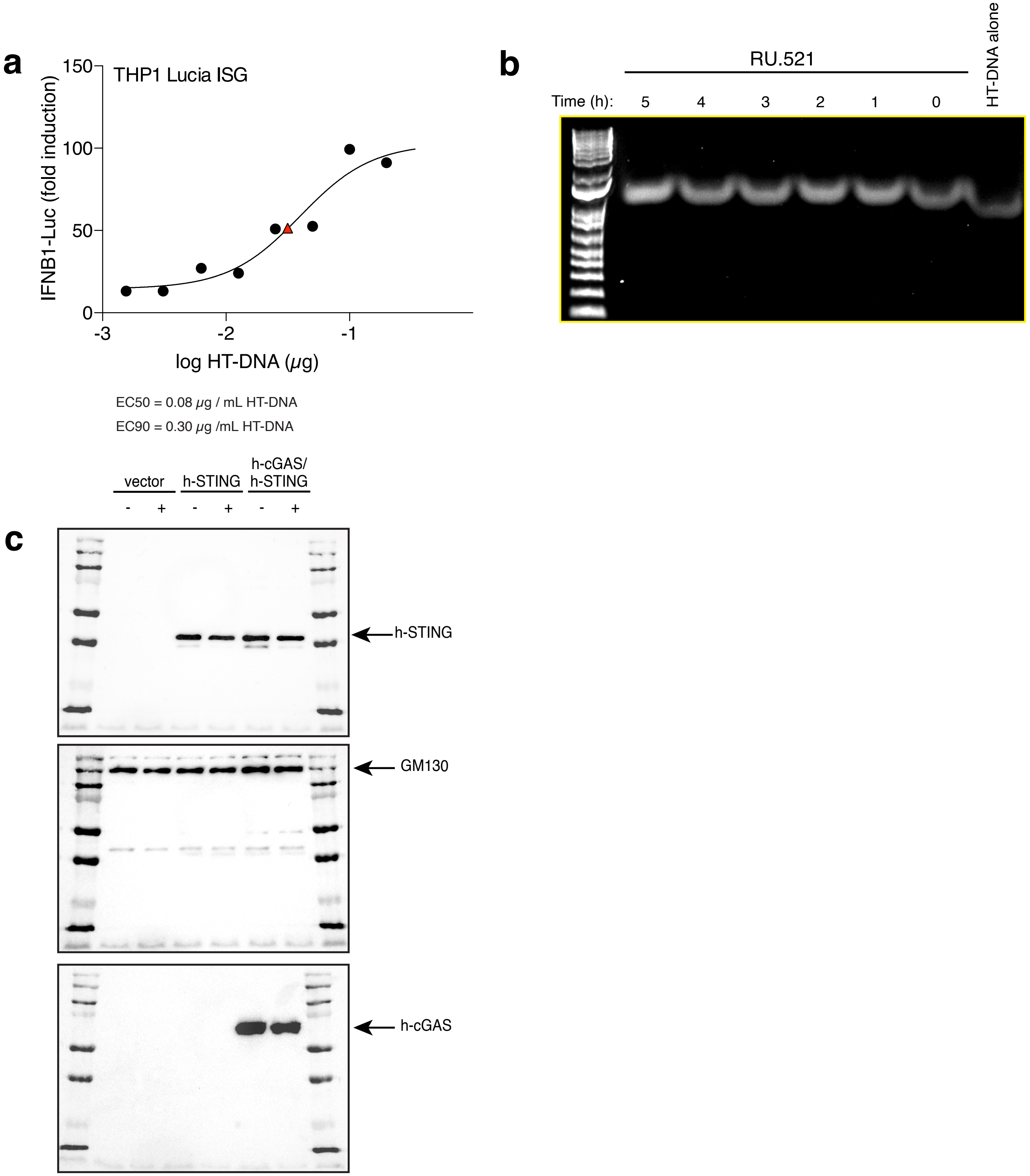


**Supplementary Fig. 1**: HT-DNA dose-response curve to determine appropriate concentration of HT-DNA for cGAS-mediated immune activation of THP-1 cells. EC90 was determined to be 0.3 µg HT-DNA/ mL of media (**a**). RU.521 does not degrade dsDNA. Using the established EC90 of 0.3 µg HT-DNA/ mL and an IC50 of 0.8 µM RU.521, both reagents were added to 1x PBS and incubated from 0-5 hours and then run on a 1.0% agarose gel (**b**). Full blot images from Figure 1 (**c**).

Supplementary Figure 2:

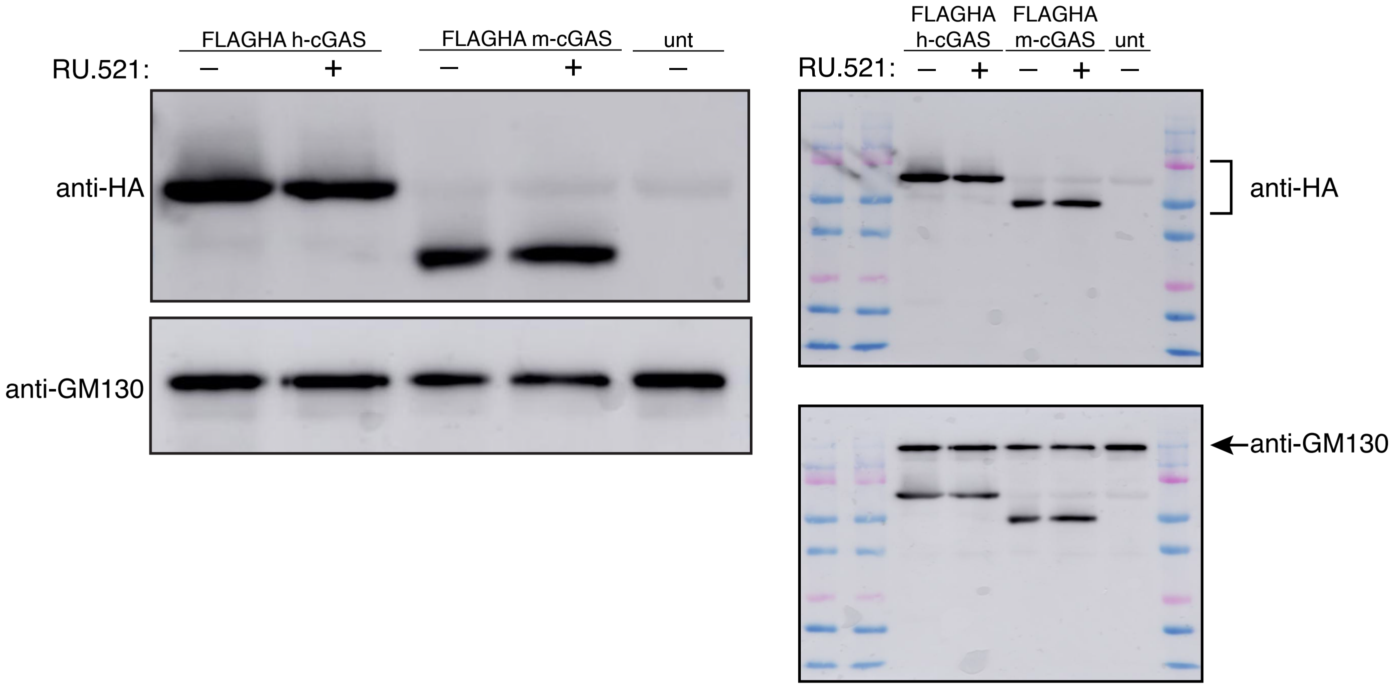


**Supplementary Fig. 2**: Western Blot of HEK293 cellular lysate transiently transfected with either human or mouse cGAS (with N-terminal Flag and HA tags). The presence of RU.521 does not alter protein levels of m-cGAS or h-cGAS. GM130 served as the loading control. 10 µg of total protein was loaded into each well.

Supplementary Figure 3:

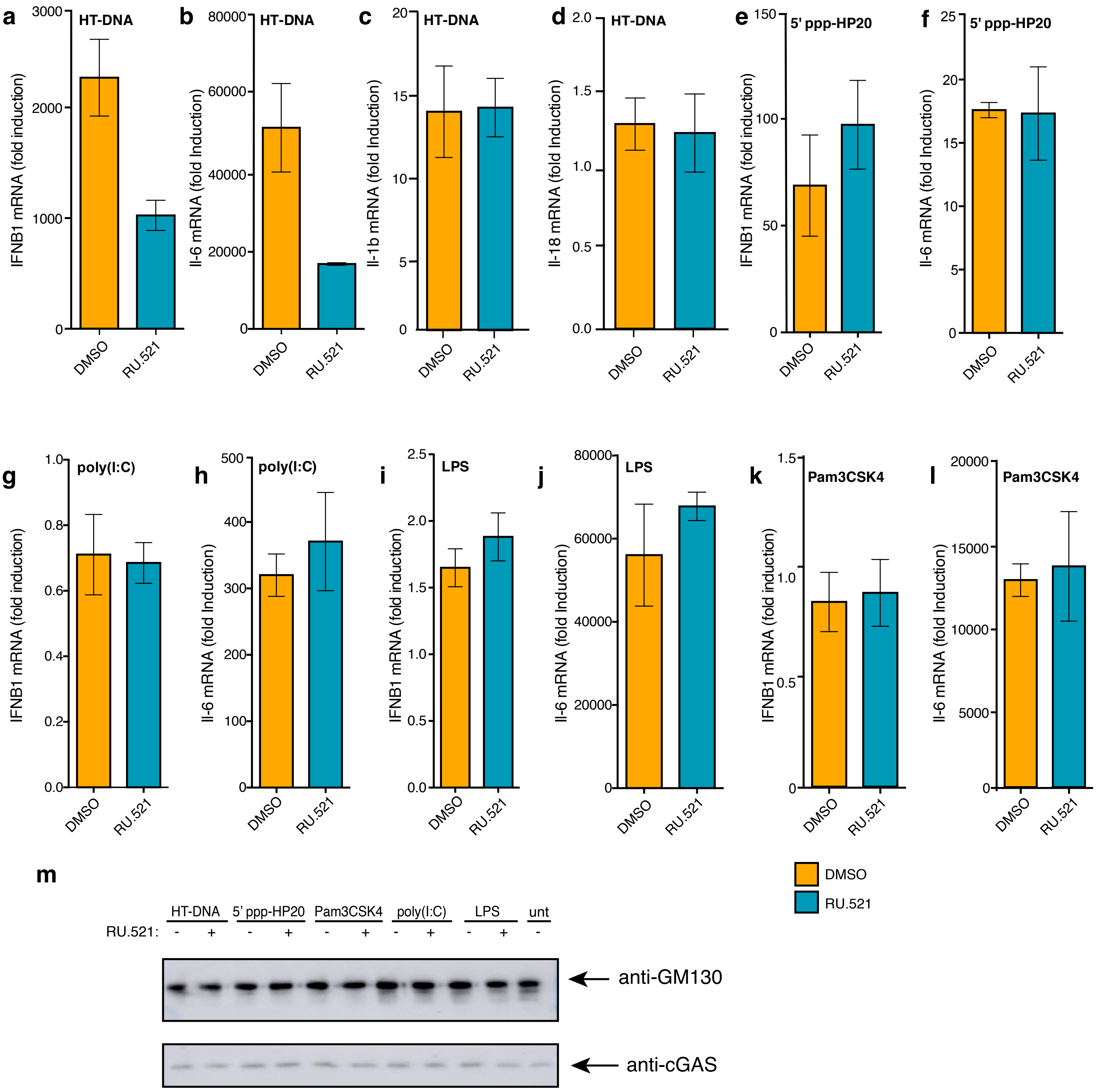


**Supplementary Fig. 3:** Actual gene expression values shown in Figure 2 (**a-l**). Immunoblot verification to ensure protein levels across samples remain consistent in the presence of RU.521 (**m**).

Supplementary Figure 4:

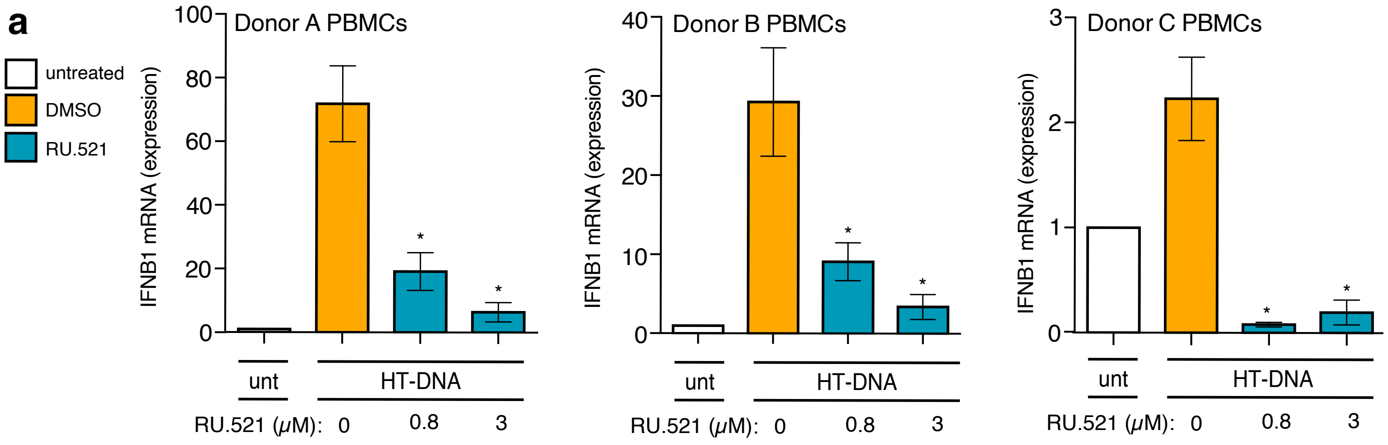


**Supplementary Fig. 4:** Actual IFNB1 mRNA expression values for each respective Donor (A,B, & C) shown in Figure 5.
